# Heterosis and Combining Ability in Pumpkin Inbreds (*Cucurbita moschata* Duch. ex Poir.)

**DOI:** 10.1101/2022.01.16.476506

**Authors:** Most. Shaika Shafin, Most. Shanaj Parvin, Md Ehsanul Haque, Fahmida Akhter

## Abstract

Twenty hybrids along with five parents evaluated in the study were mainly contemplated to find out the best cross combinations and the best general and specific combiners as well as to estimate the nature and magnitude of the gene action for different qualitative traits. Using Griffing’s and Hayman’s approach through a 5 × 5 full diallel cross fashion, an investigation on heterosis and combining ability in pumpkin was undertaken following RCBD design with three replications at the experimental field. Both positive and negative significant GCA and SCA variances were obtained from few parents and hybrids. Predominance of additive-additive gene action was noted for most of the characters except hollowness and dry matter content, where additive-dominance gene action was predominant; flesh thickness and brix (%), where dominance-dominance gene action were predominant. A single parent was not found as good combiner for more than two characters. The best specific combiners were IBD 40 X IBD 47 for beta carotene, total sugar and fruit yield; IBD23 X IBD40 for brix (%), hollowness and flesh thickness; IBD40 X IBD57 for fruit breadth; IBD47 X IBD50 for non reducing sugar; and IBD47 X IBD57 for reducing sugar. The Vr-Wr graphs exhibited complete, partial and over dominance effect of genes for different characters. Complete dominance was observed only for beta carotene whereas over dominance was noticed for hollowness and flesh thickness. Partial dominance was ensured for fruit breadth, dry matter, brix (%), reducing sugar, non-reducing sugar, total sugar and fruit yield. Significant heterosis of some crosses against mid parent and better parents were observed for some characters.

## INTRODUCTION

Pumpkin (*Cucurbita moschata* Duch. ex Poir.) is locally known as ‘Misti kumra’ or ‘Mistilau’ or ‘Misti kudu’, is an important common vegetable in Bangladesh. Pumpkin originated in equatorial and sub-equatorial America (Whitaker and Davis, 1999). It starts from Southern part of USA and continues up to Peru of South America. It grows throughout the entire tropical and sub-tropical regions of the world and milder areas of the temperate zones of hemispheres. It is widely cultivated in India, China, Malaysia, Taiwan, and Bangladesh. It is distributed widely in Southeast Asia, tropical Africa, tropical South and Central America (Peru and Mexico), the Caribbean and most part of tropics.

Pumpkin belongs to the family Cucurbitaceae. There are 27 species under the genus *Cucurbita*, five of which are in cultivation. These are *C. moschata, C. maxima, C. ficifolia, C. pepo and C. mixta*, commonly known as pumpkin. Pumpkin is highly cross pollinated crop having chromosome number 2n=40. *C. moschata* is probably the most widely grown species of *Cucurbita* and this species is cross compatible with *C. maxima, C. pepo and C. mixta*. It is insect-pollinated and 1000 m isolation distance is necessary to maintain purity of cultivars plants and vine crop. It is an annual crop having a climbing or trailing habit (Katyal and Chadha, 2000).

Pumpkin is relatively high in energy and carbohydrates and a good source of vitamins, especially high carotenoid pigments and minerals (Bose and Som, 1998). The nutrient per 100 g edible portions of fruit is cited in appendix 1. Night-blindness is a serious problem in Bangladesh which happens due to vitamin A deficiency. It may certainly contribute to improve nutritional status of the people, particularly the vulnerable groups in respect of vitamin A requirement. Encouraging the mass people to take more pumpkin can easily be solved the problem. As a matter of fact, there is a program from Health Department to encourage feeding mature pumpkin to the children.

The delicate shoots and leaves are used as delicious vegetables. The fleshy large fruits can be consumed at mature and immature stages. It is one of the main vegetable in a wedding party or on other occasional party in northern India (Chauhan, 1995). The sweet pie and pumpkin haluwa are the delicious items prepared from matured fruit (Shanmungavelu, 1998). The seed are very nutritious (it contains 40-50% oil and 30% protein) and eaten as food in many countries of the world (Tindall, 1998).

The pumpkin has good medicinal value. It is used against many diseases like gonorrhea, urinary problem. The paste of the dried fruit-stalk, which is in immediate contact with the ripe gourd, is used as the remedy for the bites of venomous insects of all kinds, especially for the centipedes (Chauhan, 1995).

It is a very common vegetable in Bangladesh and particularly popular among the rural people. It is grown round the year in our country. It becomes available even in the lean period when other vegetables are scarce in Bangladesh. It has the longest storability among all the cucurbits. The well-matured fruits (ripe fruits) can be stored for 2 to 4 months (Yawalkar, 1991). Due to its good taste and keeping quality, nutritional status, easier cooking quality, reasonable market price and year-round availability, its demand is increasing day by day in the country.

Vegetable production rate in Bangladesh is very low; yearly only 8.542 million M tones (BBS, 2010). Vegetables consumption rate is 104 g per day per adult, against the optimum amount of about 300 g per day per adult (Rashid, 2006). The total area under cultivation of pumpkin is 27,935 ha with a total production of 2, 63, 000 MT having national average of 9.42 ton/ha in a year of this country (BBS, 2012).

The productivity of local genotypes ranged from 6.93 t/ha to 19.07t/ha (Hamid *et al*., 1991). On the other hand there are many exotic genotypes, which have short life cycle but high yield potential. Some of these exotic genotypes bear deep green long fruits, which are attractive. Flower buds of these genotypes appear 20 to 25 days earlier than the local genotypes. The exotic genotypes do not need big trellis because of their medium climbing habit. However, the exotic types are more susceptible to different virus diseases than the local genotypes. These variabilities among the indigenous and exotic genotypes are genetic attributes, which can be combined through hybridization to develop short vine type varieties with high yield, with smaller fruit type with high carotene content virus resistance and high number of female flowers. Being a cross-pollinated crop, it seems easy to transfer suitable traits by crossing appropriate genotypes of sweet gourd.

Though it is a very common crop, it may be mentioned that until to date there is no released variety of pumpkin with high yield potential and better nutritional quality. Further, a very limited attempt had been made for the genetic improvement of this crop, particularly with quality traits. An understanding of the nature and magnitude of variability among the genetic stocks is of prime importance to the breeder. Good knowledge of genetic resources might also help in identifying desirable cultivars for commercial cultivation. Because of its high cross-pollination, genetically pure strain is available hardly to the growers. Lack of high yielding, disease and pest tolerant variety is the main constraint towards its production. Among the cultivated landraces, a wide range of genetic variability exists in this crop that can be exploited for its improvement. It is the touchstone to a breeder to develop high-yielding varieties through selection, either from the existing genotypes or from the segregates of a cross. Hence, information on gene action, its nature and magnitude in respect of quality characters aspects is required to be properly assessed for its improvement.

Heterosis breeding is a potential tool to achieve improvement in the quality, quantity, and productivity of pumpkins (Tamilselvi *et al*., 2015). Heterosis and combining ability is a powerful tool in identifying the best combiner that may be used in crosses either to exploit heterosis or to accumulate fixable gens and obtain desirable segregates. It will help to understand the genetic architecture of various characters that enable the breeder to design effective breeding plan for future up gradation of the existing materials. The information may also be useful to breeders for genetic improvement of the existing genotypes on the basis of performance in various hybrid combinations.

There have been not many studies on heterosis and combining ability of pumpkin in Bangladesh particularly with quality traits except Rana *et al*., (2015). Though pumpkin as a vegetable is becoming an important ingredient in daily diet, relatively little attention has been paid towards the development of hybrids/varieties which are rich in beta-carotene with high Brix content; high reducing sugar with high yielding capacity. Therefore, considering the above facts the present investigation was carried out to achieve the following objectives:

1. to identify potential parents and productive hybrids of pumpkin
2. to estimate the combining ability effects and variances for quality traits in pumpkin
3. to identify of best cross combination for higher yield and other quality characters
4. to estimate the heterosis against mid and better parents of different characters

## MATERIALS AND METHODS

### Experimental site

The experimental site is located at the centre of Modhupur tract (24.09 °N latitude and 90.26 °E longitude), which is 8.4 m above the sea level. It is about 40 km North of Dhaka, The site was previously under shal forest and developed later for research purpose.

### Climate

The Experimental site is situated in the sub tropical climate zone, characterized by heavy rainfall during the month of May to September and scanty rainfall during rest of the year. During crossing of parents, the average temperature, relative humidity and rainfall was 27.26°C (max) and 18.18 °C (min), 85.03% and 3.48 mm per month, respectively (Appendix 3). During studying combining ability of the parents and hybrids, the average temperature, relative humidity and rainfall was 28.05°C (max) and 16.28 °C (min), 78.86 % and 26.31 mm per month, respectively.

### Soil

The soil is terrace soil, which is nearly equivalent to Ochrept sub order of USDA soil taxonomy and belongs to the locally termed Salna series of Shallow Red Brown Terrace soil (Brammer, 1971). The soil is silt loam in texture having acidic (pH=5.5) in nature, poor fertility status, and impeded internal drainage.

### Materials

Five (5) advanced inbreds of pumpkin viz IBD 23, IBD 40, IBD 47, IBD 50 and IBD 57 developed by the GPB, BSMRAU was used in the study for combining ability analysis in a 5×5 diallel population. The inbreds were synthesized in the previous year.

### Design and layout

The experiment was laid out in a randomized complete block design (RCBD) with three replications.

### Raising of seedlings

The seeds were sown in 9cm x 15 cm sized polyethylene bags. Two seeds were sown in each bag. The growth medium was prepared by mixing compost and soil in 50:50 proportions. Intensive care was taken for production of healthy seedlings.

### Preparation of land and pits

The experimental land was prepared by deep and cross ploughing and harrowing followed by laddering. The plots were raised 10 cm above the ground level. Pits of 50 × 50 × 50 cm size were dug at a spacing of 2 × 2 m.

### Manure and fertilizer applied in each pit

Around 10 kg Cow dung, 52 g TSP, 60 g Urea and 40 g MP were applied in each pit. Cow dung and pit soils were mixed together. The fertilizers were applied on the top and worked up to 10 cm soil of the pits.

### Transplanting

Twenty four days old seedlings were transplanted in well-prepared experimental plot on 17^th^ December, 2013. The seedlings were watered immediately after transplanting. Four plants of each genotype were accommodated in each replicated plot maintaining 2×2 m spacing.

### Intercultural operations

Intercultural operations were done as necessary during the growing period for proper growth and development of the plants and to protect the fruits from rotting.

### Mulching

Immediately after planting, the field was covered with straw to ensure optimum moisture for easy emergence of buds.

### Weeding

Routine weeding were done to keep the field free from weeds and to pulverize the soil.

### Irrigation and drainage

Irrigation was applied as and when required.

### Harvesting

The fruits were harvested when the peduncle dried on maturity.

### Data collection

Three plants were selected at random from each plot for recording data. Both quantitative and qualitative characters were recorded.

### Quantitative characters

a. Days to first male and female flower: The number of days to first male and female flower was recorded.
b. Days to first male and female flower opening: The number of days to first male and female flower opening was also recorded.
c. Nodes for first male and female flowers: The nodes at the ground level to the nodes of first blooming of male and female flowers were recorded.
d. Number of male and female flower per plant: From the first blooming of male and female flowers were counted.
e. Number of nodes for first fruit setting: The numbers of nodes from the first fruit setting were counted.
f. Fruit yield per plant (Kg): Total numbers of fruits from three randomly selected plants were weighed and their average value was taken.
g. Fruit length: Fruit length was measured using scale.
h. Fruit breadth: Fruit breadth was measured using scale.
i. Hollowness: Fruit hollowness was measured using scale after cutting the whole fruit into two pieces.
j. Flesh Thickness: Flesh thickness was measured using scale.

### Qualitative Characters

a. Dry matter (%): Dry matter percentage was also calculated from matured fruit. For dry matter content, 200gm of matured pumpkin was cut into small pieces and dried in the sun for 3 to 4 days. After that it was again kept in an oven at 60°c for 72 hours. Then the weight was taken using electric balance.
b. Brix (%): It was measured with the help of a Brix meter (Model: ATAGONI Brix 0-32%, Made in Japan).
c. Carotene (mg/g): Three fruits of each genotype were used for carotene analysis and their average value was taken. At first 10 g flesh of pumpkin was taken and crushed by mortar and pastel. After then 10 ml mixture (Acetone: Hexane=2:3) was added in the paste. Then the solution of pumpkin paste and acetone hexane was filtered in a vial having air tight lid. Then the nascent spectrophotometer reading was recorded at four different nano meter length viz 663 nm, 645 nm, 505 nm and 453 nm. Finally, β-carotene was calculated by the following formula: B-carotene (mg) = (Reading of 663 nm) +(Reading of 453 nm)-(Reading of 645 nm) - (Reading of 505 nm).
d. Sugar (gm/100gm) Estimation

One fruits of each genotype from each replication were used for reducing sugar analysis and their average value was taken.

### Reducing Sugar

Ten ml of each of Bertrand A (40g of CuSO_4_. 5H_2_O dissolved in water and diluted to 1 liter) and Bertrand B (200g of sodium-potassium tartarate and 150g of NaOH dissolved in water and diluted to 1 liter) solutions were added to 5ml of sample solution. The conical flask was placed on a hot plate (sand bath) and boiled for about 3 minutes and kept overnight for cooling. The supernatant was decanted and discarded very carefully by keeping the precipitation. The precipitation was washed repeatedly until blue color was present. Then 10ml of Bertrand C [50g of Fe_2_ (SO_4_)_3_ and 115ml of concentrated H_2_SO_4_ was added and diluted to 1 litre] solution was added to dissolve the precipitation (Cu_2_O). Finally, it was titrated with 0.4% KMnO_4_ solution. Reducing sugar was calculated comparing tabulated values. Before calculation of reducing sugar factor of 0.4% KMnO_4_ was determined.

### Total Sugar

Ten ml of extract solution was taken in a volumetric flask and 2-3 drops of 4N HCl was added. The flask was then boiled for about 3 minutes on a hot plate for hydrolysis. After cooling in tap water, the extract was neutralized with 0.1N NaOH. The rest of the procedure was same as mentioned in reducing sugar.

### Non-Reducing Sugar

Non-reducing sugar was calculated by deducting reducing sugar from total sugar.

### Statistical Analysis

Method 1 model II of Griffing (1956b) was followed for combining ability analysis. The recorded data were analyzed by using Diallel analysis and simulation program by Mark, D. Burow and James G. Coors, copyright 1993 and version 1.1.Vr-Wr graph of different traits was drawn according to Hayman (1954).

## RESULTS AND DISCUSSION

### Analysis of variance (ANOVA) for heterosis and combining ability

The mean sum of squares from analysis of variance due to heterosis and combining ability for fruit length, fruit breadth, hollowness, flesh thickness, dry matter (%), brix (%), reducing sugar, non reducing sugar, total sugar and fruit yield have been shown in Table 1.

**Table 1:** Analysis of variance (ANOVA) of combining ability for different characters in a five parental full diallel population Of pumpkin.

| Source of Variation | df | Mean sum of square |  |  |  |  |  |  |  |  |  |  |
| --- | --- | --- | --- | --- | --- | --- | --- | --- | --- | --- | --- | --- |
|  |  | FL | FB | HN | FT | DRM | BRX | BCAR | RS | NRS | TS | FY |
| Replication | 2 | 1.11** | 6.82** | 0.22 | 0.03 | 4.480* | 0.001 | 0.133 | 0.130 | 0.001 | 0.173 | 7.983** |
| Genotype | 24 | 1.07** | 5.23** | 0.68 | 0.36 | 8.670** | 0.016 | 1.262 | 0.368 | 2.11** | 5.136** | 9.997** |
| GCA | 4 | 1.57** | 4.34** | 1.32 | 0.10 | 1.060** | 0.015 | 1.921 | 0.735 | 4.35** | 5.206** | 7.891** |
| SCA | 10 | 6.73** | 3.63** | 0.34 | 0.55 | 5.465** | 0.018 | 1.019 | 0.229 | 0.968 | 6.628** | 1.023** |
| Reciprocal | 10 | 1.21** | 7.19** | 0.75 | 0.27 | 1.110** | 0.015 | 1.242 | 0.359 | 2.360* | 3.616* | 1.343** |
| Error | 48 | 2.078 | 1.068 | 0.120 | 0.412 | 1.021 | 0.001 | 0.170 | 0.095 | 0.217 | 0.381 | 6.269 |
| GCA/SCA |  | 2.330 | 1.159 | 3.816 | 0.180 | 1.940 | 0.833 | 1.885 | 3.209 | 4.498 | 0.785 | 0.077 |

FL= Fruit length, FB= Fruit breadth, HN= Hollowness, FT=Flesh thickness, DRM= Dry matter (%), BRX=Brix%, BCAR= Beta carotene (mg/100g), RS= Reducing sugar (g/100 gm), NRS=Non reducing sugar (g/100g), TS=Total sugar (g/100g), FY=Fruit yield (kg/plant)

Analyses of variance for genotypes were highly significant for fruit length, fruit breadth, hollowness, brix (%) and fruit yield. The analysis of variance for general combining ability variances were found highly significant for the traits fruit length, fruit breadth, hollowness, brix (%), dry matter percentage and fruit yield (Table 1) which indicated that additive gene actions played significant role for the expression of these characters.

General combining ability was found non-significant for the character such as flesh thickness, β carotene, reducing sugar, non reducing sugar and total sugar. Specific combining ability and reciprocal variances were found significant for fruit length; fruit breadth, hollowness, brix (%) and fruit yield which indicated that non additive gene action were involved in the inheritance of these traits. The characters viz. flesh thickness, β carotene, reducing sugar, non reducing sugar and total sugar showed non-significant specific combining ability and reciprocal variation.

The ratio of GCA and SCA variance was high and more than one for the characters fruit length, hollowness, dry matter, β carotene and reducing sugar revealed that the preponderance of additive gene action over the non-additive gene action. This indicated the limited scope of heterosis breeding for these characters and population improvement through recurrent selection should be adopted for exploiting the genetic variations (Kushwaha and Ram, 1996; Pandey *et al*., 2005, Jha *et al*., 2009). The gca variances was lower than the sca variance for fruit breadth, flesh thickness, brix (%), non reducing sugar, total sugar and fruit yield, may be improved through hybridization (heterosis) as indicating the predominance of non-additive gene effects which was opined by Jha *et al*. (2009). However both additive and non-addictive gene actions were observed by Suneal Kumar (1984) and Sirohi *et al*., (1986).

### Mean performance of parents

The characters studied of the parents were fruit length, fruit breadth, hollowness, flesh thickness; dry matter (%), brix (%), reducing sugar, non-reducing sugar, total sugar and fruit yield and their mean values are presented in Table 2.

**Table 2:**
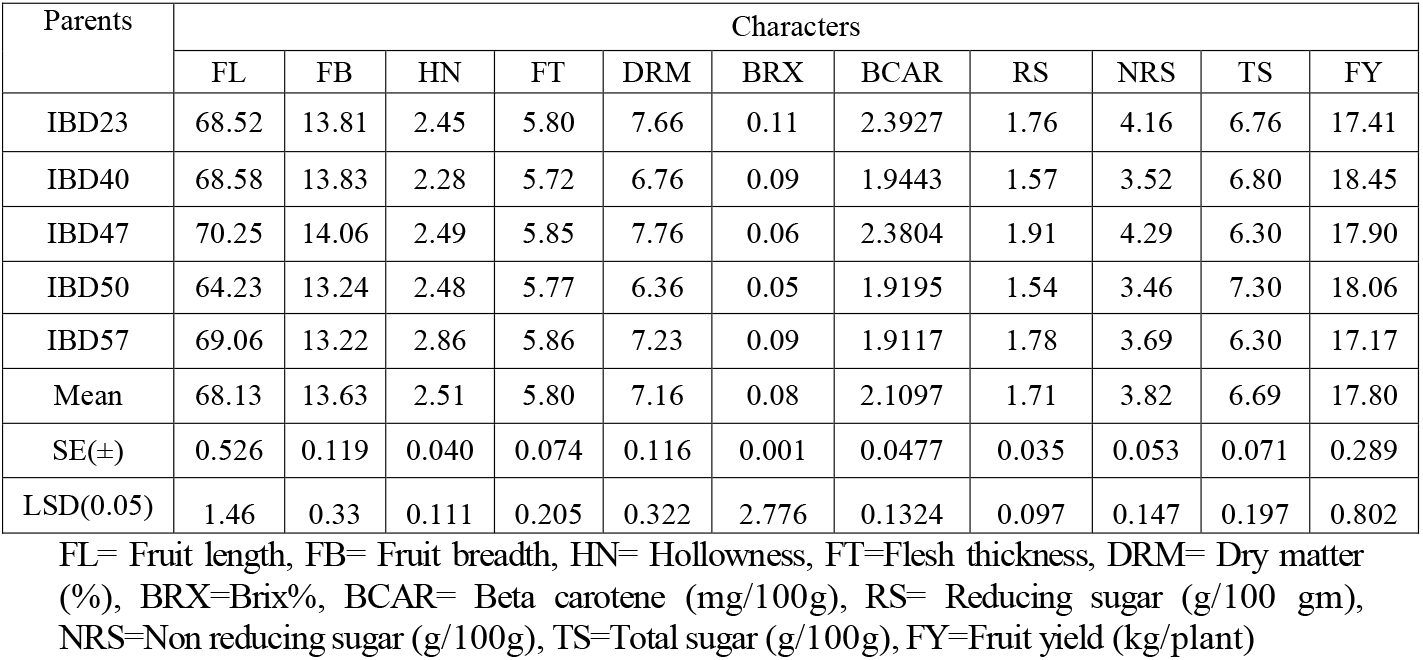
Mean performance of five parents for different quantitative and qualitative characters in Pumpkin.

Significant variations among the parents for different traits were recorded. Variation among the parents for fruit length was observed. The highest fruit length was ascertained in IBD 47 and the lowest in IBD 50 having value (64.23cm)_;_ other three parents had fruit length more or less similar to parental mean (68.72 cm). The highest fruit breadth was observed in IBD 47 which was statistically similar to IBD23, IBD40, IBD50 and IBD57. Hollowness was found the highest in case of IBD57; and the lowest in IBD40. The highest flesh thickness was found in IBD23, IBD47 and IBD57; other two parents having the lowest value were noticed in IBD40 and IBD50. In case of dry matter content, the highest value was observed in IBD47 and the lowest value was ascertained from IBD50. Higher brix (%) was found in the parent IBD23 and lower value was found in IBD50. IBD40 and IBD57 possessed similar value. Beta carotene content was the maximum in case of IBD23 and IBD57; other three parents had beta carotene content more or less similar to parental mean (1.924 %). Non-reducing sugar and reducing sugar was the highest in case of IBD47 but total sugar was higher in case of IBD 50. Reducing sugar and non reducing sugar was the lowest in parent IBD50. In case of total sugar parent IBD47 and IBD 57 contained lower value. The highest mean value for fruit yield was found in IBD40 and the lowest in case of IBD57.

### Mean performance of hybrids

Significant variations among the hybrids for different traits were observed. This study now proceeds to observe the cross-breeding population for conducting genetic analysis of mean values of F_1_s (Table 3) exposed that fruit length was the highest in the cross combination IBD57 X IBD50 and the lowest was found in cross combination IBD50 X IBD40.

**Table 3:** Mean performance of 20 crosses for different quantitative and qualitative characters in pumpkin.

| Crosses | Characters |  |  |  |  |  |  |  |  |  |  |
| --- | --- | --- | --- | --- | --- | --- | --- | --- | --- | --- | --- |
|  | FL | FB | HN | FT | DRM | BRX | BCAR | RS | NRS | TS | FY |
| IBD23 X IBD40 | 63.70 | 13.21 | 2.20 | 6.27 | 8.33 | 8.8 | 0.2440 | 1.62 | 4.06 | 7.00 | 13.93 |
| IBD23 X IBD47 | 65.00 | 13.00 | 1.62 | 5.80 | 7.00 | 6.6 | 0.2055 | 1.92 | 3.98 | 4.00 | 7.50 |
| IBD23 X IBD50 | 64.70 | 12.91 | 2.26 | 5.24 | 6.33 | 6.4 | 0.2197 | 1.99 | 4.19 | 7.00 | 15.50 |
| IBD23 X IBD57 | 66.78 | 13.18 | 2.70 | 5.20 | 6.66 | 6.4 | 0.2517 | 2.08 | 4.60 | 7.66 | 17.10 |
| IBD40 X IBD23 | 76.00 | 15.91 | 2.53 | 6.34 | 7.33 | 7.6 | 0.1762 | 1.59 | 3.36 | 7.00 | 24.81 |
| IBD40 X IBD47 | 75.00 | 16.00 | 2.31 | 5.98 | 6.66 | 6.4 | 0.2058 | 1.85 | 3.91 | 9.33 | 29.14 |
| IBD40 X IBD50 | 65.83 | 14.25 | 2.30 | 5.35 | 6.00 | 5.4 | 0.1127 | 1.28 | 2.40 | 8.00 | 21.68 |
| IBD40 X IBD57 | 74.33 | 12.96 | 2.31 | 5.61 | 7.00 | 6.4 | 0.1501 | 1.81 | 3.31 | 7.00 | 20.30 |
| IBD47 X IBD23 | 78.10 | 14.45 | 3.43 | 5.73 | 12.00 | 11.8 | 0.3020 | 2.15 | 5.17 | 4.66 | 16.03 |
| IBD47 X IBD40 | 68.41 | 13.33 | 2.16 | 5.82 | 7.66 | 6.6 | 0.2605 | 1.48 | 4.09 | 7.00 | 20.55 |
| IBD47 X IBD50 | 68.31 | 14.25 | 2.36 | 6.60 | 7.33 | 6.8 | 0.4198 | 1.81 | 6.01 | 7.00 | 17.28 |
| IBD47 X IBD57 | 71.83 | 14.66 | 2.93 | 5.75 | 9.00 | 8.6 | 0.1801 | 1.79 | 3.59 | 7.00 | 24.11 |
| IBD50 X IBD23 | 61.10 | 12.50 | 1.96 | 5.63 | 7.00 | 6.4 | 0.1740 | 1.28 | 3.02 | 8.33 | 16.26 |
| IBD50 X IBD40 | 55.11 | 10.15 | 1.83 | 5.58 | 5.66 | 5.4 | 0.1446 | 1.44 | 2.89 | 6.00 | 11.18 |
| IBD50 X IBD47 | 67.10 | 14.16 | 2.40 | 5.64 | 5.00 | 5.0 | 0.2026 | 1.23 | 3.25 | 7.33 | 22.08 |
| IBD50 X IBD57 | 63.66 | 13.00 | 3.40 | 6.22 | 5.10 | 5.0 | 0.1333 | 1.43 | 2.76 | 5.66 | 12.35 |
| IBD57 X IBD23 | 72.91 | 13.16 | 3.50 | 5.97 | 6.66 | 6.4 | 0.2096 | 1.82 | 3.92 | 5.33 | 17.06 |
| IBD57 X IBD40 | 70.66 | 14.33 | 2.66 | 5.58 | 6.33 | 6.4 | 0.2148 | 1.58 | 3.73 | 5.33 | 15.06 |
| IBD57 X IBD47 | 64.83 | 12.00 | 2.35 | 5.98 | 8.33 | 8.6 | 0.3010 | 2.79 | 5.10 | 6.66 | 14.13 |
| IBD57 X IBD50 | 79.75 | 15.35 | 2.93 | 5.96 | 11.33 | 11.8 | 0.2184 | 1.88 | 4.07 | 7.66 | 31.58 |
| Mean | 65.23 | 13.63 | 2.50 | 5.81 | 7.33 | 6.8 | 0.2127 | 1.74 | 3.87 | 7.09 | 18.38 |
| SE | 2.63 | 0.59 | 0.20 | 0.37 | 0.58 | 0.008 | 0.2382 | 0.17 | 0.26 | 0.35 | 1.44 |
| LSD (0.05) | 5.50 | 1.23 | 0.41 | 0.77 | 1.21 | 0.016 | 0.4981 | 0.35 | 0.54 | 0.73 | 3.01 |

FL= Fruit length, FB= Fruit breadth, HN= Hollowness, FT=Flesh thickness, DRM= Dry matter (%), BRX=Brix%, BCAR= Beta carotene (mg/100g), RS= Reducing sugar (g/100g), NRS=Non reducing sugar (g/100 gm), TS=Total sugar (g/100g), FY=Fruit yield (kg/plant)

For the trait fruit breadth, the highest and the lowest mean value was observed in the combination IBD40 X IBD47 and IBD50 X IBD40, respectively. Hollowness exhibited the highest value in IBD57 X IBD23. Flesh thickness was found the highest in cross IBD40 X IBD57 and the lowest was found in the cross IBD23 X IBD57. The highest dry matter percentage was found in IBD47 X IBD23 and lowest in IBD50 X IBD47. The highest and the lowest brix (%) were observed in IBD40 X IBD57and IBD47 X IBD40, respectively. For the trait beta carotene, higher value was found in the cross IBD47 X IBD50 and the lowest in the combination IBD40 X IBD50. The highest total sugar content was noticed in IBD40 X IBD47 whereas the highest reducing and non reducing sugar was ascertained from the cross combination IBD57 X IBD47 and IBD47 X IBD50, respectively. For the trait fruit yield, the highest mean value was observed in the cross combination IBD57 X IBD50 and the lowest was in IBD23 X IBD47.

### General combining ability (GCA) effect

The information regarding GCA effect of the parent is of prime importance as it is difficult to pickup good general combiner for all the characters (Jha *et al*., 2009). A parent with higher positive significant GCA effects is considered as a good general combiner. The magnitude and direction of the significant effects for different parents provide meaningful comparison and would give a route to the future breeding program. The GCA effect of five parents for eleven different characters along with standard error and standard error of difference are presented in the Table (4a, 4b). No single parent contained all the desirable characteristics. Moreover, these parents could be utilized in the crossing program depending on the objectives.

**Table 4a:**
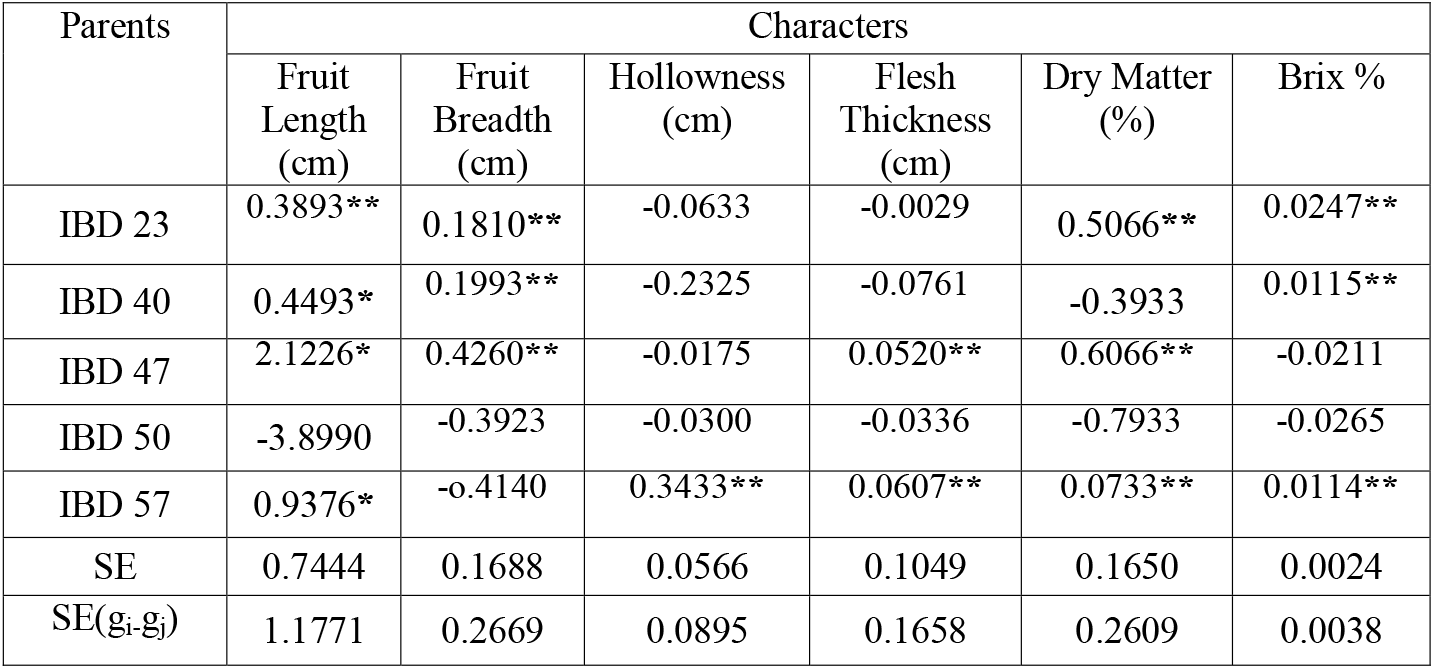
GCA effects for fruit length, fruit breadth, hollowness, flesh thickness, dry matter and brix (%) in 5 × 5 full diallel populations in pumpkin.

**Table 4b:** GCA effects for β carotene, reducing sugar, non reducing sugar, total sugar and fruit yield in 5 × 5 full diallel populations in pumpkin.

| Parents | Characters |  |  |  |  |
| --- | --- | --- | --- | --- | --- |
| | $\beta$ Carotene (mg/100g) | Reducing Sugar (g/100gm) | Non Reducing Sugar (g/100g) | Total Sugar (g/100g) | Fruit Yield (kg/plant) |
| IBD 23 | 0.283** | 0.051** | 0.334** | 0.073** | -0.392 |
| IBD 40 | -0.165 | -0.141 | -0.307 | 0.106** | 0.653** |
| IBD 47 | 0.270** | 0.201** | 0.472** | -0.393 | 0.103** |
| IBD 50 | -0.190 | -0.175 | -0.365 | 0.606** | 0.262** |
| IBD 57 | -0.198 | 0.063** | -0.134 | -0.393 | -0.626 |
| SE(±) | 0.067 | 0.050 | 0.076 | 0.100 | 0.408 |
| SE(gi-gj) | 0.106 | 0.079 | 0.120 | 0.159 | 0.646 |

### Fruit length

The estimates of GCA effects for fruit length are given in the Table 4a. Positive GCA effect is preferable to increase the fruit size. All the parents except IBD50 had significant GCA effect. The parent IBD23 provided the highest positive significant GCA effects (0.389**) followed by IBD57 (0.938**\***). These two parents could be selected as good combiner for increased fruit length. The other parent IBD50 had negative significant GCA effects which could be selected as good combiner for reduced fruit length. Sirohi and Choudhury (1977) also found significant GCA effects for fruit length in a 8 × 8 half diallel experiments in bitter gourd.

### Fruit breadth

The estimates of GCA effects for this character are given in the Table 4a. Positive GCA effect is preferable to increase the fruit breadth and negative GCA effects to reduce the fruit breadth. The three parents viz. IBD23, IBD40 and IBD47 had positive significant effects for this trait. So these three parents could be selected as a good combiner for increasing fruit breadth. Rana *et al*. (2015) found similar results.

### Hollowness

Cavity hollowness is inversely proportional to flesh thickness. The GCA effects for all the parents had non-significant (Table 4a) value except IBD57. The parent IBD57 provided positive significant GCA effects (0.343**). Thus this parent could be a combiner for this trait, though negative significant value as observed in other four parents which expected to reduce the hollowness.

### Flesh Thickness

The highest positive GCA effects (Table 4a) was found in IBD47 (0.052**) followed by IBD57 (0.061**), which was significant as positive significant value is expected to increase the flesh thickness. The GCA effects of other three parents were non-significant. So, the three parents viz. ca IBD23, IBD40, and IBD50 were not suitable for the improvement of this character.

### Dry matter (%)

The data on GCA effects for dry matter are given in Table 4a. The positive significant effects was provided by the parent IBD23 (0.530**), IBD47 (0.606**) and IBD57 (0.073**) which is expected as it indicates to increase the dry matter percentage. Among the five parents, IBD40 and IBD50 demonstrated non-significant GCA effects, so these two could not be used a good combiner for future breeding program of this character.

### Brix (%)

Brix index is a parameter which measures the flesh sweetness of a variety in pumpkin. GCA effects for this trait was found significant (Table 4a) for the three parents i.e. IBD23 (0.025**), IBD40 (0.011**) and IBD57 (0.011**) which would serve as a good combiner for increasing the sweetness of flesh.

### β carotene

The GCA effects for this character were found significant for two parents (Table 4b). The parent IBD23 provided the highest positive significant GCA effects (0.243**\*\***) followed by IBD47 (0.271**\*\***). The GCA effects were non-significant for other three parents (IBD40, IBD50 and IBD57). Rana *et al*., (2015) and Pandey *et al*., (2010) found similar results in pumpkin.

### Reducing Sugar

The positive and significant GCA effects (Table 4b) on reducing sugar obtained by the parent IBD23 (0.052**), IBD47 (0.201**) and IBD57 (0.064**) which could be used for further breeding program to increase reducing sugar content in pumpkin. The two other parents (IBD40 and IBD50) had non-significant GCA effects i.e. these parents would not be suitable for improvement of reducing sugar in pumpkin.

### Non reducing sugar

The counts on GCA effects for non-reducing sugar are given in Table 4b. The highest positive significant GCA effects was found in parent IBD47 (0.472**) followed by IBD23 (0.335**) demonstrated that these two parents could be selected for increasing non-reducing sugar content in pumpkin. Among all other parents, IBD40, IBD50 and IBD57 had non-significant effects.

### Total Sugar

The positive and significant GCA effects (Table 4b) was found in IBD50 (0.606**), IBD40 (0.106**) and IBD23 (0.073**) for this trait i.e. the rest three parents were not suitable for improvement of total sugar content in pumpkin.Yang *et al*. (2006) also found similar results in pumkin.

### Fruit yield

Significant GCA effects were ascertained from three parents for fruit yield (Table 4b) but other two parents had negative value. From GCA effects analysis it implied that among five parents, IBD40 (0.653**), IBD47 (0.103**), and IBD50 (0.262**) might be selected as a good combiner for increasing fruit yield per plant. Nisha and Veeraragavathatham (2014) had similar results in their experiments.

### Specific combining ability (SCA) effect and reciprocal effect

Non-additive gene action is signified by the SCA effects in the expression of the characters of a crop. Specific combining ability indicated the performance of some specific cross combination. That is why it is related to a particular cross. High SCA effects may arise not only in crosses involving high general combiners but also in those involving low combiners. Estimates on SCA effects of the crosses in F_1_ generation revealed that there were a good number of crosses having significant positive or negative SCA effects on different important traits of pumpkin. The SCA effect of 10 F_1_ for eleven different characters studied are presented in the Table 5a, 6a. Reciprocal effects are presented in the Table 5b, 6b.

**Table 5a:** Specific combining ability (SCA) effects for fruit length, fruit breadth, hollowness, flesh thickness and dry matter in 5 × 5 full diallel population of pumpkin.

| Crosses | SCA Effects |  |  |  |  |
| --- | --- | --- | --- | --- | --- |
|  | FL | FB | HN | FT | DRM |
| IBD23 X IBD 40 | 0.880 | 0.552** | 0.150** | 0.587** | 0.560** |
| IBD 23 X IBD47 | 0.907 | -0.516 | 0.093** | -0.083 | 1.226** |
| IBD 23 X IBD50 | -1.721** | -0.714 | -0.306 | -0.329 | -0.206 |
| IBD 23 X IBD57 | 0.392 | -0.226 | 0.303** | -0.272 | -1.073 |
| IBD 40 X IBD47 | 1.005 | 0.407** | -0.025 | 0.125** | -0.206 |
| IBD 40 X IBD50 | -4.206** | -1.241 | -0.187 | -0.226 | -0.140 |
| IBD 40 X IBD57 | 2.982 | 0.230** | -0.135 | -0.191 | -0.173 |
| IBD47 X IBD50 | 1.354 | 0.540** | -0.085 | 0.302** | -0.806 |
| IBD 47 X IBD57 | -2.857** | -0.312 | -0.200 | -0.045 | 0.826** |
| IBD50 X IBD57 | 6.539 | 1.347** | 0.336** | 0.265** | 1.726** |
| SE(sij) | 1.534 | 0.348 | 0.116 | 0.216 | 0.340 |
| SE(sij-skl) | 2.038 | 0.462 | 0.155 | 0.287 | 0.452 |

FL= Fruit length, FB= Fruit breadth, HN= Hollowness, FT=Flesh thickness, DRM= Dry matter

**Table 5b:**
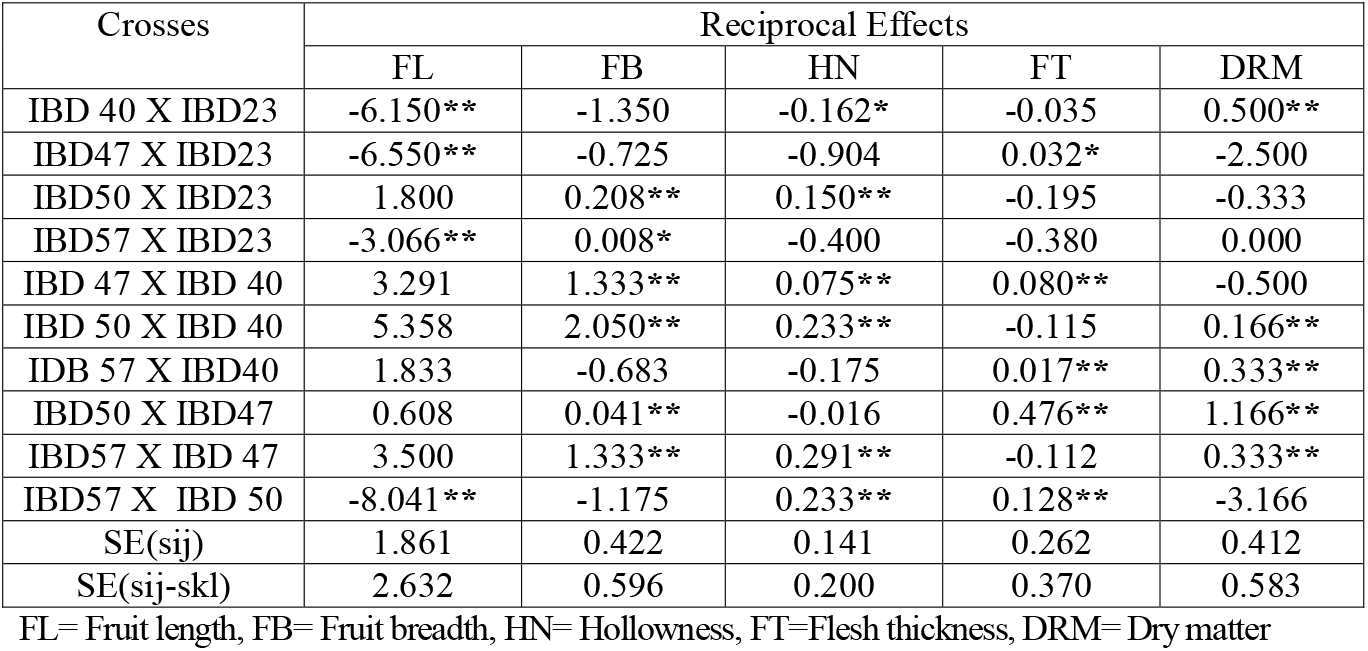
Reciprocal effects for fruit length, fruit breadth, hollowness, flesh thickness and dry matter in a 5 × 5 full diallel populations of pumpkin.

**Table 6a:** Specific combining ability (SCA) effects for brix(%), beta carotene, reducing sugar, non-reducing sugar, total sugar and fruit yield in a 5 × 5 full diallel populations of pumpkin.

| Crosses | SCA Effects |  |  |  |  |  |
| --- | --- | --- | --- | --- | --- | --- |
|  | BRX | BCAR | RS | NRS | TS | FY |
| IBD23 X IBD40 | -0.028 | -0.126 | -0.013 | -0.139 | 0.126** | 1.310* |
| IBD23X IBD50 | 0.012** | -0.125 | 0.068** | -0.057 | -2.042 | -5.748* |
| IBD23 X IBD47 | 0.002** | -0.233 | 0.047** | -0.185 | 0.293** | -1.790* |
| IBD23 X IBD57 | -0.031 | 0.112** | 0.122** | 0.234** | 0.126** | 0.297* |
| IBD40 X IBD47 | -0.067 | 0.116** | -0.103 | 0.012** | 1.760** | 6.245* |
| IBD40 X IBD50 | -0.030 | -0.467 | -0.035 | -0.503 | -0.406 | -2.286* |
| IBD40 X IBD57 | 0.121** | 0.078** | 0.060** | 0.139** | -0.240 | -0.148* |
| IBD47 X IBD50 | 0.034** | 0.922** | -0.221 | 0.700** | 0.260** | 1.513* |
| IBD47 X IBD57 | -0.040 | -0.131 | 0.313** | 0.182** | 0.926** | 1.843* |
| IBD50 X IBD57 | -0.045 | 0.037** | 0.053** | 0.091** | -0.240 | 4.526* |
| SE(sij) | 0.005 | 0.139 | 0.103 | 0.156 | 0.207 | 0.842 |
| SE(sij-sk1) | 0.006 | 0.184 | 0.138 | 0.208 | 0.276 | 1.119 |

BRX=Brix, RS= Reducing sugar, NRS=Non reducing sugar, TS=Total sugar, FY=Fruit yield, BCAR=Beta carotene

**Table 6b:** Reciprocal effect for brix (%), beta carotene, reducing sugar, non reducing sugar, total sugar and fruit yield in 5 × 5 full diallel populations of pumpkin.

| Crosses | Reciprocal Effects |  |  |  |  |  |
| --- | --- | --- | --- | --- | --- | --- |
|  | BRX | BCAR | RS | NRS | TS | FY |
| IBD40 X IBD 23 | -0.064 | 0.338** | 0.014** | 0.353** | 0.000 | -5.441** |
| IBD47 X IBD23 | -0.058 | -0.482 | -0.112 | -0.595 | -0.333 | -4.266** |
| IBD50 X IBD23 | -0.065 | 0.228* | 0.355* | 0.583** | -0.666 | -0.383** |
| IBD57 X IBD23 | 0.069* | 0.210** | 0.128** | 0.338** | 1.166** | 0.016 |
| IBD47 X IBD40 | -0.002 | -0.273 | 0.184** | -0.089 | 1.166** | 4.295* |
| IBD50 X IBD40 | -0.016 | -0.159 | -0.082** | -0.242 | 1.000** | 5.250 |
| IBD 57 X IBD40 | 0.076** | -0.323 | 0.115** | -0.207 | 0.833** | 2.616 |
| IBD 50 X IBD47 | -0.054 | 1.085** | 0.291** | 1.377** | -0.166 | -2.400** |
| IBD 57 X IBD47 | -0.008 | -0.249 | -0.503 | -0.753 | 0.166** | 4.991 |
| IBD57 X IBD50 | -0.007 | -0.425 | -0.226 | -0.651 | -1.000 | -9.616** |
| SE(sij) | 0.006 | 0.168 | 0.126 | 0.190 | 0.252 | 1.022 |
| SE(sij-skl) | 0.008 | 0.238 | 0.178 | 0.269 | 0.356 | 1.445 |

BRX=Brix, RS=reducing sugar, NRS=Non reducing sugar, TS=Total sugar, FY=Fruit yield, BCAR= Beta carotene

The SCA values provide important information about the performance of the hybrid relative to its parents. However, Arunga *et al*. (2010) found that the SCA effect alone has limited value for parental choice in breeding programs. They, therefore, suggested that the SCA effects should be used in combination with other parameters, such as hybrid mean and the GCA of the respective parents such that a hybrid combination with both high mean and favorable SCA estimates and involving at least one of the parents with high GCA, would tend to increase the concentration of favorable alleles; which is desired by any breeder (Tamilselvi *et al*., 2015).

### Fruit length

Out of 10 F_1_s, all hybrids provided non-significant positive or negative SCA effects (Table 5a) on fruit length indicating their unsuitability to increase or decrease fruit length.

Out of 10 F_1_s, all cross combination showed reciprocal effect. IBD57 X IBD23 provided the highest negative and significant reciprocal effects (Table 5b) on fruit length followed by IBD40 X IBD23 and IBD47 X IBD23, having value **-**3.066 **, -6.150** and **-**6.550**, respectively indicated their suitability of using maternal effect to increase fruit length (fruit size). On the other hand, the cross IBD50 X IBD40 (5.358) gave the highest positive reciprocal effects followed by IBD57 X IBD47, IBD47 X IBD40 and IBD57 X IBD40 having value (3.500, 3.291and 1.833), respectively which can be used to reduce fruit size to some extent. Monhanty (2000) and Rana *et al*., (2015) agreed with our findings.

### Fruit breadth

The cross combination IBD50 X IBD57 exhibited the highest significant and positive SCA effects (1.34**) followed by IBD23 X IBD40 (0.552**). Thus these cross could be considered as good specific combiner for increasing fruit breadth in pumpkin. The negative SCA value (Table 5a) for this parameter was obtained from the crosses viz., IBD23 X IBD 57 and IBD47 X IBD57 could provide decreased fruit breadth in pumpkin.

Among all ten crosses, all crosses had reciprocal effects of which six crosses had significant and positive reciprocal effects (Table 5b) indicated their ineptness (unsuitability) to increase fruit breadth in pumpkin.

### Hollowness

Among the cross combination, four crosses viz. IBD23 X IBD40 (0.150**), IBD23 X IBD47 (0.093**), IBD23 X IBD57 (0.303**) and IBD50 X IBD57 (0.336**) showed significant positive SCA values (Table 5a). The rest six the cross combinations which had negative values were non-significant for hollowness in pumpkin.

The data about reciprocal effects for hollowness are given in Table 5b. All the crosses showed reciprocal effects. Among the cross, IBD57 X IBD47(0.291**) had the highest positive and significant reciprocal effects on hollowness of pumpkin followed by IBD57 X IBD 50, IBD50 X IBD40, IBD50 X IBD23 and IBD47 X IBD40 having value 0.233**,0.233**, 0.150**and 0.075**, respectively. Hence these cross combinations could provide chance to reduce the hollowness in pumpkin which is expected. On the other hand, the hybrid IBD40 X IBD23 (−0.162**\***) gave the negative significant reciprocal effects. Similar results were also found by Pandey *et al*., (2010) and Rana *et al*., (2015).

### Flesh thickness

Positive and significant value is expected in case of flesh thickness (Table 5a) in order to increase flesh thickness and the yield as well. The cross combination IBD23 X IBD40 revealed the highest (0.476**) positive and significant SCA value which could be used as a good specific combiner for flesh thickness in pumpkin.

The cross combination IBD50 X IBD47 (0.476**) had the highest positive and significant reciprocal effects on of flesh thickness of pumpkin followed by IBD57 X IBD50, IBD47 X IBD23, IDB57 X IBD40, and IBD47 X IBD40 having value 0.128**,0.032**\***,0.017** and 0.080**, respectively which could be used as the good specific combination to increase the flesh thickness in pumpkin.

### Dry matter (%)

The estimates on SCA effects for dry matter are given in (Table 5a).The highest positive and significant effect was found in cross IBD50 X IBD57(1.726**) followed by IBD23 X IBD47 (1.226**), IBD47 X IDB57 (0.826**) and IBD23 X IBD40(0.560**), respectively which is expected as it indicated increase in dry matter percentage. All other cross combination had non-significant and negative SCA effects for this trait.

The estimates on reciprocal effects for dry matter are given in Table 5b.Among the cross IBD50 X IBD47 (1.166**) had the highest positive and significant reciprocal effects on dry matter (%) of pumpkin followed by IBD40 X IBD23 and IBD50 X IBD40 with values 0.500** and 0.166**, respectively. The cross combinations IDB57 X IBD40 and IBD57 X IBD 47 revealed similar positive and significant reciprocal effects (0.333**) which could be specific combination to increase the dry matter (%) in pumpkin. Among all ten crosses, all crosses had reciprocal effects except IBD57 X IBD23. Rana *et al*., (2015) and Tamilselvi *et al*., (2015) observed similar trend in their experiment in pumpkin.

### Brix (%)

Among the 10 crosses, four revealed significant SCA effects and six showed non-significant SCA effects (Table 6a). The combination IBD40 X IBD57 exhibited the highest (0.121**\*\*)** positive and significant SCA effects followed by IBD47 X IBD50, IBD47 X IBD50 and IBD23 X IBD50 with SCA values 0.034**, 0.012** and 0.002**, respectively. Hence these combinations could be regarded as a good specific combiner to increase brix (%) in pumpkin.

Out of 10, only two crosses revealed significant reciprocal effects and the rest eight showed non-significant reciprocal effects (Table 6b). The cross combinations IBD57 X IBD40 and IBD57 X IBD23 exhibited positive and significant reciprocal effects with 0.076** and 0.069**\***, respectively. Therefore these combinations could also be used as specific combiner to increase brix (%) in pumpkin.

### β carotene

The SCA effects for this character was significant and positive for the combinations IBD23 X IBD57, IBD40 X IBD47, IBD40 X IBD57, IBD47 X IBD50 and IBD50 X IBD57 (Table 6a). To increase β carotene content in pumpkin, it is necessary to have positive significant value. The estimates on reciprocal effects for beta carotene are given in (Table 6b).Among the cross, IBD50 X IBD47 (1.085**) had the highest positive and significant reciprocal effects on β carotene content in of pumpkin followed by IBD40 X IBD23, IBD50 X IBD23 and IBD57 X IBD23, having values 0.338**, 0.228**\*** and 0.210**, respectively.

### Reducing Sugar

Reducing sugar indicated the sweetness of a variety. The highest positive and significant SCA effects (Table 6a) on reducing sugar obtained from the cross combination IBD47 X IBD57 (0.313**) followed by IBD23 X IBD57 (0.122**), IBD23 X IBD47 (0.068**), IBD50 X IBD57 (0.053**) and IBD23 X IBD50 (0.047**) which could be used for further breeding program to increase reducing sugar content in pumpkin. The rest combinations had negative and non-significant SCA effect. There existed reciprocal effects in the crosses. The highest negative and significant reciprocal effects (Table 6b) on reducing sugar obtained from the cross combination IBD50 X IBD23 (0.355**\***) followed by IBD50 X IBD47, IBD47 X IBD40, IBD57 X IBD23, IBD57 X IBD40 and IBD40 X IBD23 with values 0.291**, 0.184**, 0.128**, 0.115**and 0.014**, respectively which could be used for further breeding program to increase reducing sugar content in pumpkin.

### Non reducing sugar

The counts on SCA effects for non reducing sugar are given in (Table 6a). Among the 10 crosses six revealed positive and significant SCA effects and four showed negative and non-significant SCA effects. The highest positive and significant SCA effects on non-reducing sugar obtained from the cross combination IBD47 X IBD50 (0.700**) followed by IBD23 X IBD57 (0.234**), IBD47 X IBD57 (0.182**), IBD40 X IBD57 (0.139**), IBD50 X IBD57 (0.091**) and IBD40 X IBD47 (0.012**).

The data on reciprocal effects for non reducing sugar are given in (Table 6b). The parents showed reciprocal effects in specific crosses. The highest and positive significant SCA effect was found in the cross combination IBD50 X IBD47 (1.377**) followed by IBD50 X IBD23, IBD40 X IBD23 and IBD57 X IBD23 with values 0.583**, 0.353** and 0.338**, respectively which demonstrated that these combination could be selected to increase non reducing sugar content in pumpkin. Rest of all combinations had negative and non-significant SCA effect.

### Total Sugar

The highest and positive significant SCA effects (Table 6a) was found in cross IBD40 X IBD47 (1.760**\*\*)** followed by IBD47 X IBD57, IBD23 X IBD50, IBD47 X IBD50 and IBD23 X IBD57 with SCA values of 0.926**, 0.293**, 0.260** and 1.760**, respectively for this trait. The cross combinations IBD23 X IBD40 and IBD23 X IBD57 revealed similar positive and significant effects (0.126**\*\*)** which could be specific combination to increase the total sugar content in pumpkin. To increase total sugar content it is expected to have positive and significant effects. There existed reciprocal effects for total sugar. The highest positive and significant reciprocal effects (Table 6b) was found in cross combinations IBD57 X IBD23 and IBD47 X IBD40 with similar value of 1.16** for this trait followed by IBD50 X IBD40 (1.00**), IBD57 X IBD40(0.833**) and IBD57 X IBD47(0.166**), respectively. Among all ten crosses, all crosses had reciprocal effects except IBD40 X IBD23.

### Fruit yield

For fruit yield significant and positive SCA effects was ascertained from the cross IBD40 X IBD47 with a value of 6.245**\***(Table 6a). All of ten cross combinations were significant but four crosses were negatively significant which reduced the fruit yield. Hence the positive and significant combinations could be selected for increasing fruit yield per plant in pumpkin.

The significant and negative reciprocal value (Table 6b) for this parameter was obtained from the crosses viz., IBD57 X IBD50, IBD40 X IBD23, IBD47 X IBD23, IBD50 X IBD47 and IBD50 X IBD23 which could provide decreased fruit yield per plant in pumpkin. The cross combination **I**BD47 X IBD40 demonstrated the highest significant and positive reciprocal effects (4.295**\***). Hence the positive and significant cross combination IBD50 X IBD40, IBD57 X IBD47, IBD57 X IBD40 and IBD57 X IBD23, could be used for further breeding program to increase fruit yield per plant in pumpkin.

### Vr-Wr graph and Wr+Vr/parental mean graph

#### Analysis of variance in Hayman’s analysis (following Morley Jones)

Analysis of variance showed that the significant values of ‘a’ for the character fruit length, hollowness, brix (%), beta carotene, reducing sugar and non reducing sugar suggested that additive components were involved in the regulation of these characters. The dominance component (b) was highly significant indicated that this component was important in genetic control of most of the character studied except fruit length, dry matter, reducing sugar and fruit yield. (Table 7). Item ‘b_1_’ was highly significant for two characters-flesh thickness and beta carotene, which detected unidirectional dominance and significant difference between parental and hybrid grand mean for these two characters. An asymmetrical distribution of dominant genes was suggested by the significant ‘b_2_’ value for the characters beta carotene, non reducing sugar and total sugar. The ‘b_3_’values were also significant for most of the character studied except fruit length, dry matter, reducing sugar and fruit yield which indicated the dominance deviations which are not attributable to ‘b_1_’ and ‘b_2_’and showed important contribution to the non-additive gene action. Significant ‘c’ value for the trait fruit length, brix (%), beta carotene, reducing sugar and total sugar indicated the presence of maternal effect. Significant’d’ value indicated presence of reciprocal difference in the character viz. fruit length, hollowness, brix (%), reducing sugar and non reducing sugar.

**Table 7:** Analysis of variance (ANOVA) in Hayman’s analysis (following Morley Jones) for different characters in a five parental full diallel populations of pumpkin.

| Source of Variation | df | Mean sum of square |  |  |  |  |  |  |  |  |  |  |
| --- | --- | --- | --- | --- | --- | --- | --- | --- | --- | --- | --- | --- |
|  |  | FL | FB | HN | FT | DRM | BRX | BCAR | RS | NRS | TS | FY |
| a | 4 | 1.01* | 2.092* | 6.681* | 1.224** | 0.41** | 8.05 | 1.547 | 1.381** | 1.42** | 1.001 | 1.66** |
| b<sub>1</sub> | 1 | 3.288 | 3.896* | 1.536 | 7.06* | 2.667 | 5.878** | 0.35* | 4.04 | 2.10** | 8.47 | 7.34 |
| b<sub>2</sub> | 4 | 8.3* | 6.027** | 3.95** | 1.45 | 1.43 | 1.059 | 1.36 | 1.78** | 1.43 | 1.79* | 4.74 |
| b<sub>3</sub> | 5 | 4.526 | 2.578** | 1.052 | 4.712** | 2.036* | 2.965** | 9.03 | 3.565 | 2.759 | 9.488 | 1.75** |
| b | 10 | 2.299 | 1.330 | 5.430** | 2.485 | 1.050 | 1.583 | 5.09** | 1.894** | 1.458 | 4.900** | 9.69* |
| c | 4 | 9.56* | 7.517 | 3.114** | 1.257** | 5.573** | 9.013 | 5.51** | 1.263** | 9.22** | 2.993** | 5.94 |
| d | 6 | 0.63 | 4.342** | 1.890* | 7.64** | 3.586** | 5.513** | 1.48 | 7.56* | 5.38* | 1.893** | 3.56** |
| Error | 48 | 9.558 | 9.977 | 1.398 | 5.768 | 1.981 | 4.904 | 0.010 | 8.216 | 4.575 | 1.042 | 3.257 |

FL= Fruit length, FB= Fruit breadth, HN= Hollowness, FT=Flesh thickness, DRM= Dry matter (%), BRX=Brix%, BCAR= Beta carotene (mg/100g), RS= Reducing sugar (g/100g), NRS=Non reducing sugar (g/100 gm), TS=Total sugar (g/100g), FY=Fruit yield (kg/plant)

The analysis of variance due to diallel progenies indicated significant differences among themselves, which warrants further analysis. Hayman’s graphic approach to diallel analysis is based on monogenic additive model. In this approach Hayman’s graphical analysis (Vr-Wr graph) was done and the findings are presented as Vr-Wr graphs, the two dimensional depiction made based on the parental variance (Vr) and parent offspring co-variance (Wr) which are presented in the Fig.(1-11, a) for eleven characters studied.

### Fruit length

The regression of Wr on Vr for fruit length (Fig. 1a) gave a slope b = 0.248 ± 0.126 which was less than 1.0 indicating presence of additive-additive nature of genetic system. The regression line intersected above the point of origin which indicated the presence of partial dominance for fruit length. The distribution of array points in the graph suggested that the parental genotypes P_2_, P_5_, P_3_ and P1 apparently contained frequency of dominant alleles while P_4_ had the most recessive alleles as it fall far away to the point of origin.

**Figure 1:**
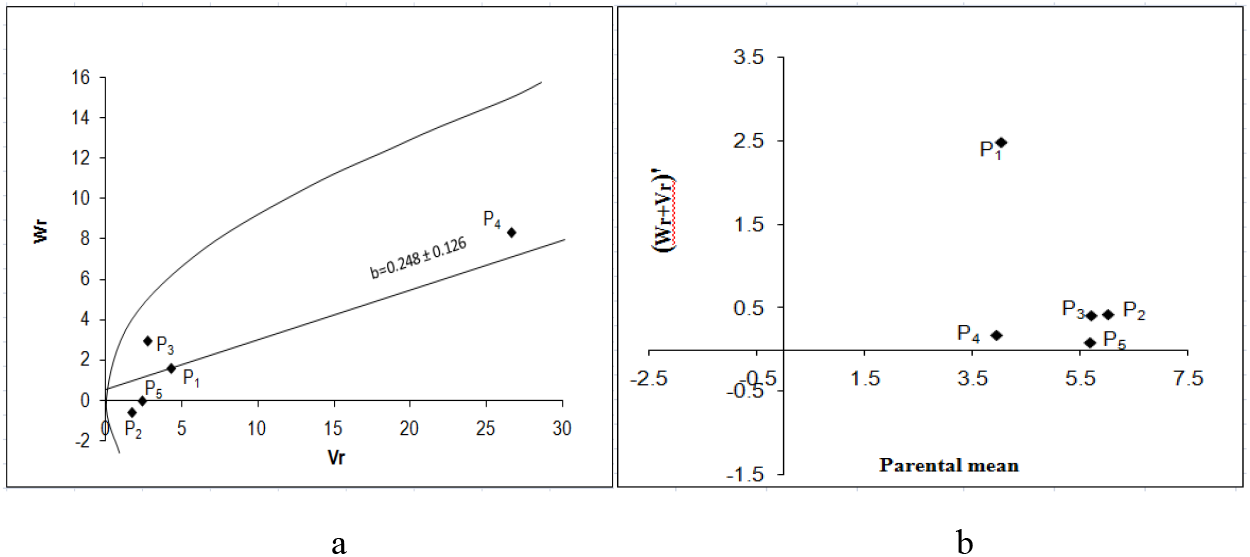
Vr - Wr graphs (a) and Wr+Vr/parental mean graph (b) for fruit length

It was observed from the Wr+Vr/parental mean graph (Fig. 1b) that all the parents possess recessive alleles associated with positive effect would result increased fruit length. Contrarily, dominant alleles will have negative effect.

### Fruit breadth

The Vr - Wr graph (Fig. 2a) for fruit breadth gave a slope b = -0.452 ± 0.426 which was negative indicating presence of non allelic interaction i.e. epistasis playing role for this trait. The regression line intersected above the point of origin which indicated the presence of partial dominance for fruit breadth. The distribution of array points indicated that among five parents P_1_ contained a frequency of dominant alleles and P_5_ possessing the maximum recessive alleles and other parents are intermediate between two. Array points scattered all along the regression line in this graph indicated genetic diversity among parents.

**Figure 2:**
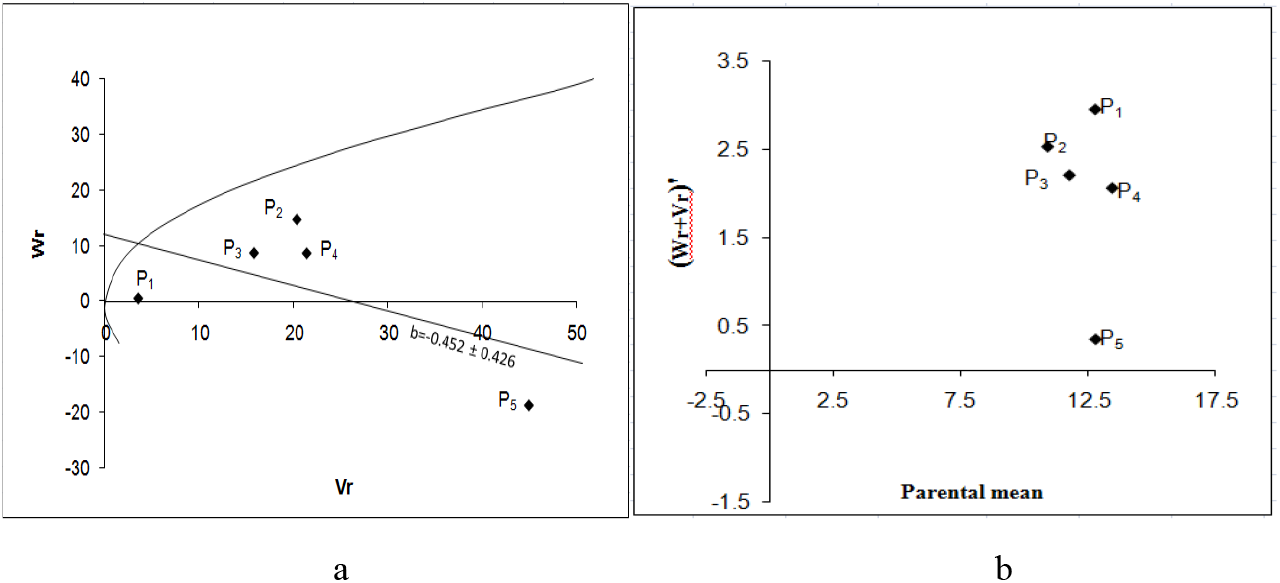
Vr – Wr graphs (a) and Wr+Vr/parental mean graph (b) for fruit breadth

The Wr+Vr/parental mean graph further confirmed the consistency of dominance against the parental score and the parental mean for this trait suggested that all the parents contained recessive alleles and most of the parents were high scoring (Fig. 2b). So, parents having higher fruit breadth were consistently associated with recessive alleles in the direction of higher value.

### Hollowness

The Vr - Wr graph (Fig. 3a) for hollowness of fruit that the regression line intersected above the point of origin which indicated the presence of partial dominance for fruit hollowness. The distribution of array points in the graph suggested that the parent P_3_ contained a frequency of dominant alleles and all other parents contained a frequency of recessive alleles. The regression of Wr on Vr gives the slope b = -0.038±0.440 which was negative indicating presence of non allelic interaction i.e. epistasis playing role for this trait.

**Figure 3:**
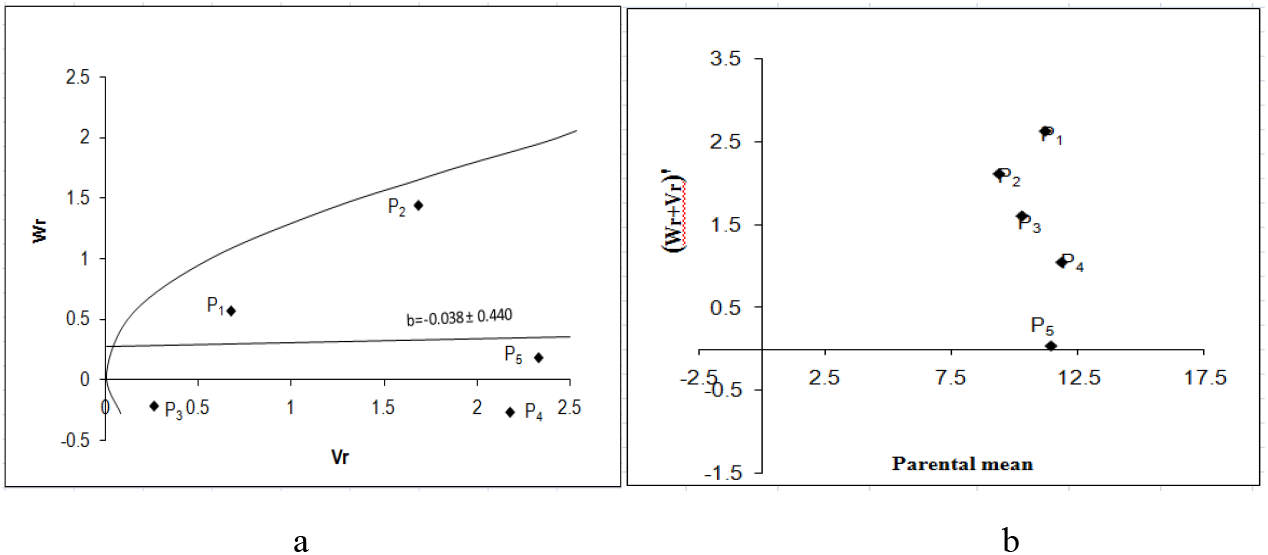
Vr - Wr graphs (a) and Wr+Vr/parental mean graph (b) for hollowness.

The Wr+Vr versus parental mean graph confirmed that all the five parents possesssed recessive alleles associated with positive effect i.e high hollowness of fruit (Fig. 3b).

### Flesh thickness

The regression of Wr on Vr for flesh thickness (Fig. 4a) gave a slope b = 0.390 ± 0.378 which was significantly far away from 1.0 indicating presence of additive-dominance nature of genetic system. The regression line intersected below the point of origin suggested over dominance gene action for controlling the trait. The distribution of array points indicated that among five parents P_2_ contained the maximum frequency of dominant alleles as it held the closest position to the point of origin. The parent P_4_ contained the maximum frequency of recessive alleles. Array points sprinkled all along the regression line in this graph indicated genetic diversity among parents.

**Figure 4:**
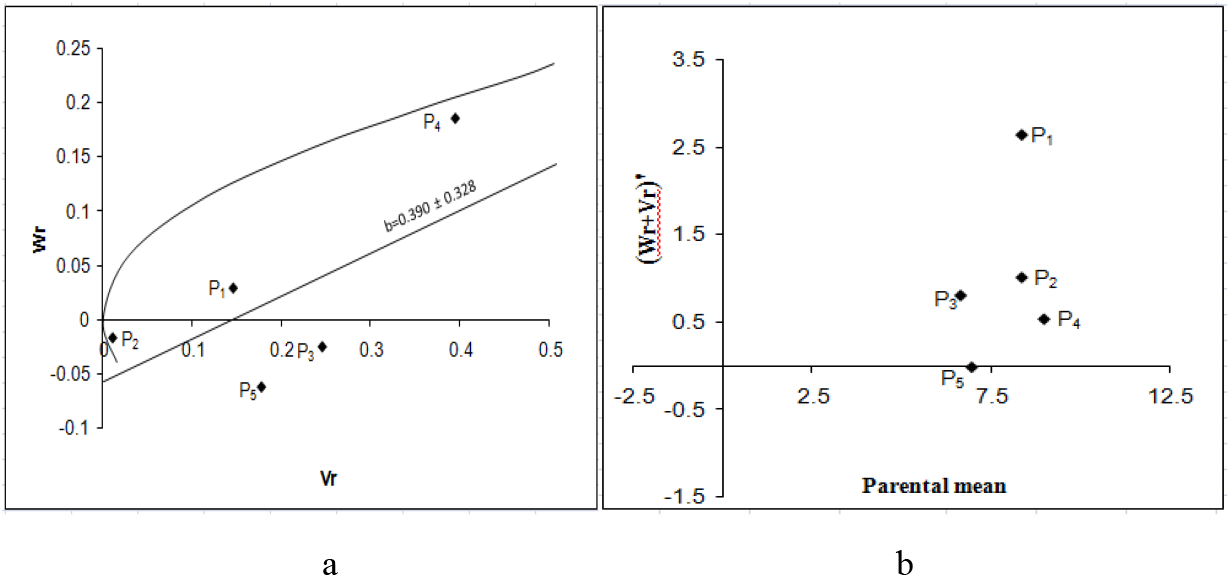
Vr - Wr graphs (a) and Wr+Vr/parental mean graph (b) for flesh thickness

The Wr+Vr/parental mean graph (Fig. 4b) tested the consistency of dominance against parental score. Parental mean suggested that all the parents contained the recessive alleles. Among them, P_4_ had the highest value as it held on the top most position, all other had moderate score. So, higher flesh thickness was associated with recessive alleles in the direction of higher values.

### Dry matter (%)

The Vr - Wr graph (Fig. 5a) for dry matter showed that the regression line intersected above the point of origin which indicated the presence of partial dominance for dry matter percentage. The distribution of array points in the graph suggested that the parent P_5_ occupying the closest position to the origin possessed the maximum frequency of dominant alleles and the parent P_1_ contained the maximum frequency of recessive alleles. Array points sprinkled all along the regression line in this graph indicated genetic diversity among parents. The regression of Wr on Vr gives the slope b = -0.582±0.445 which was negative affirming presence of non allelic interaction i.e. epistasis playing role for this trait.

**Figure 5:**
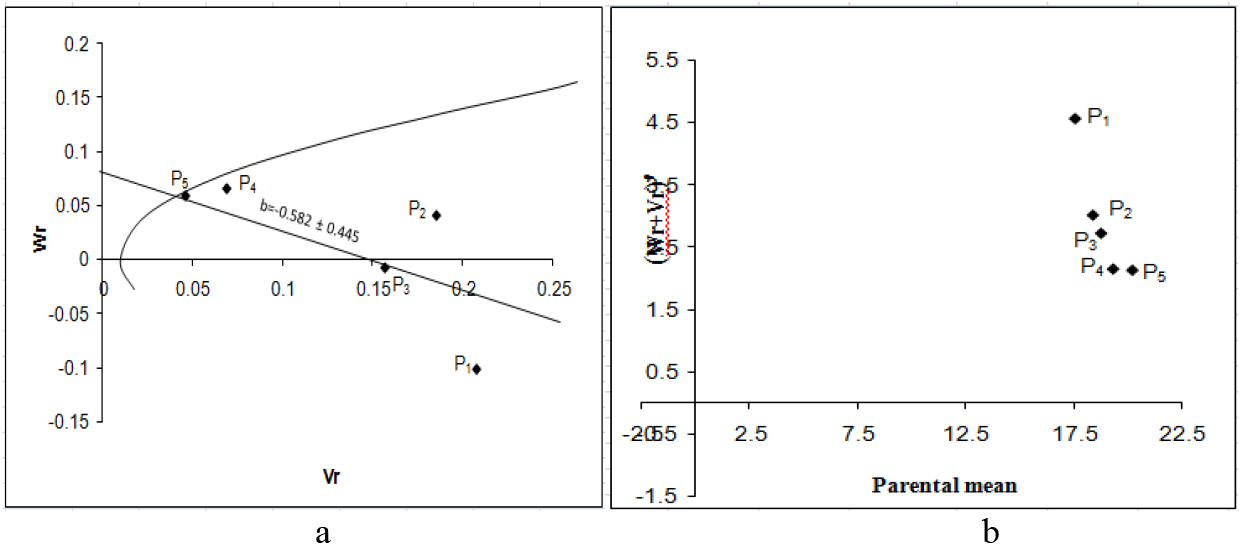
Vr - Wr graphs (a) and Wr+Vr/parental mean graph (b) for dry matter percent

The Wr+Vr versus parental mean graph (Fig. 5b) tested the consistency of dominance against parental score. Parental mean suggested that all the parents contained recessive alleles. The parent P_3_ possessed recessive alleles were high scoring whereas parents P_5_ with recessive alleles were low scoring for this trait. High dry matter content was therefore associated with parents having recessive alleles.

### Brix %

The Vr - Wr graph (Fig. 6a) for brix (%) showed that the regression line intersected above the point of origin which indicated the presence of partial dominance for brix (%). The distribution of array points in the graph suggested that the parental genotypes P_2_ and P_1_ apparently contained the large number of dominant alleles, while P_5_ had the most recessive alleles. Dispersed array points all along the regression line in this graph indicated genetic diversity among parents. The regression of Wr on Vr gives the slope b = -0.150±0.231 which was negative affirming presence of non-allelic interaction i.e. epistasis playing role for this trait.

**Figure 6:**
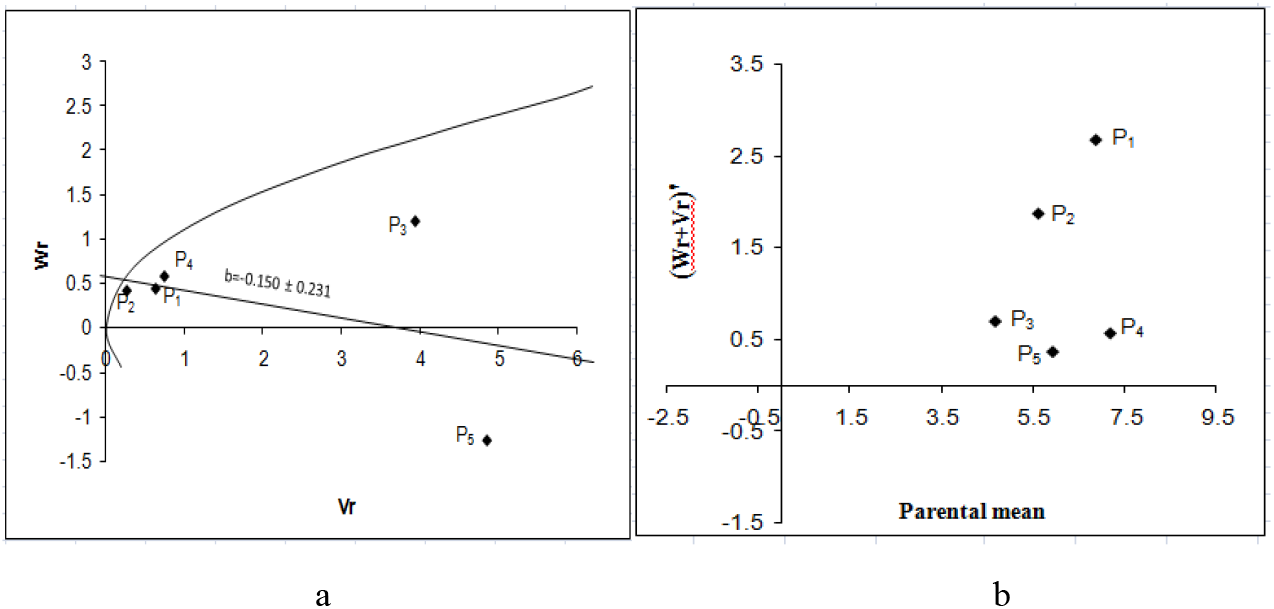
Vr - Wr graphs (a) and Wr+Vr/parental mean graph (b) for brix (%)

The Wr+Vr versus parental mean graph (Fig. 6b) affirmed that the brix (%) was conditioned by recessive alleles were high scoring for the parent P_5_ while parent P_1_ had the lowest score. All the parental values were positive Therefore; parents having high brix (%) were consistently associated with recessive alleles in the direction of higher value.

### β carotene

The regression of Wr on Vr for beta carotene (Fig. 7a) gave a slope b = 0.041 ± 0.117 which was less than 1.0 indicating presence of additive-additive nature of genetic system. The regression line intersected above the point of origin suggesting partial dominance gene action for controlling the trait. The distribution of array points indicated that among five parents P_2_ and P_5_ contained the maximum frequency of dominant alleles. The maximum frequency of recessive alleles was found in P_4_ as it falls far away to the point of origin. Array points sprinkled all along the regression line in this graph indicated genetic diversity among parents.

**Figure 7:**
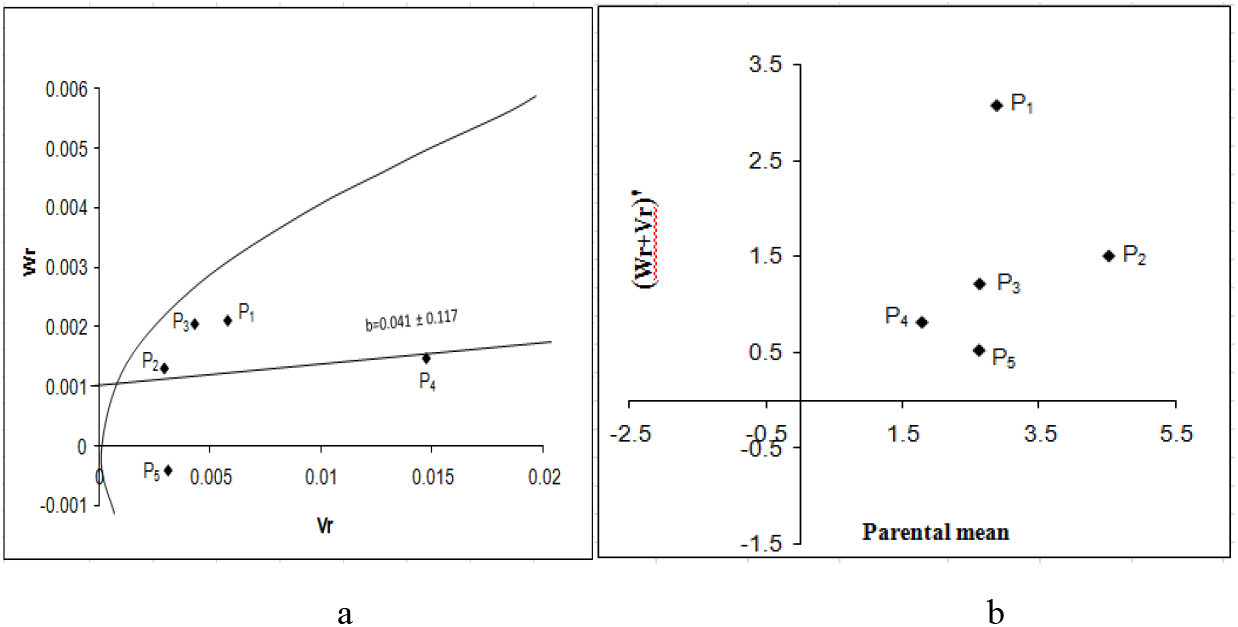
Vr - Wr graphs (a) and Wr+Vr/parental mean graph (b) for beta carotene

The Wr+Vr/parental mean graph (Fig. 10b) tested the consistency of dominance against parental score. All the five parents possessed recessive alleles associated with positive effect for β carotene content.

**Figure 8:**
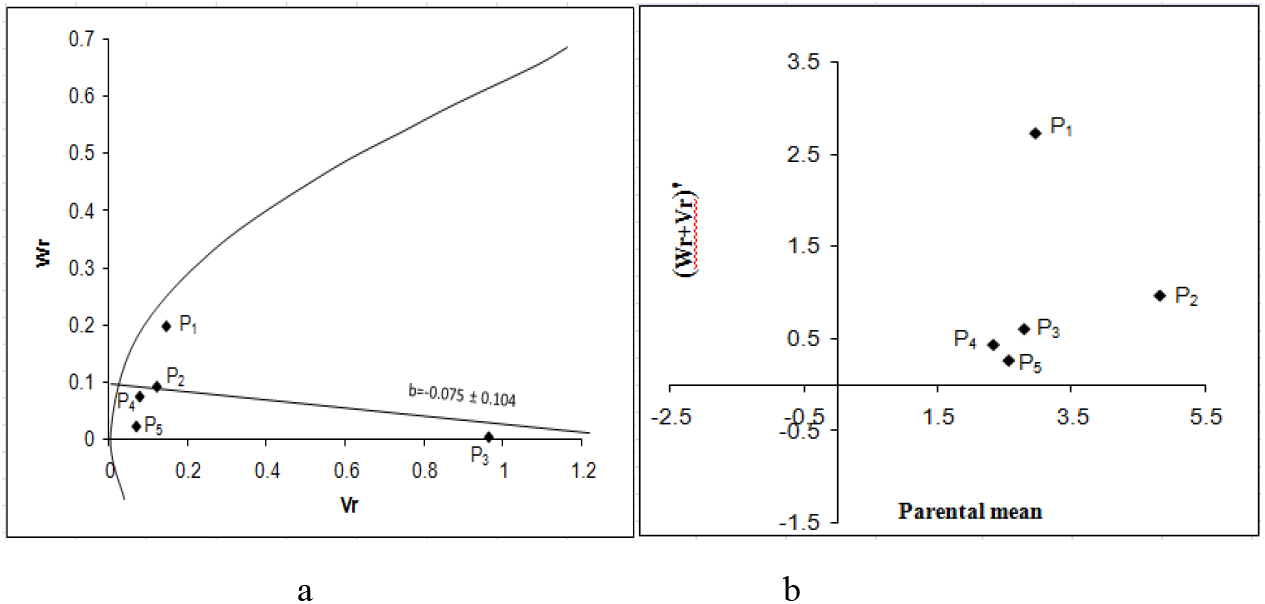
Vr - Wr graphs (a) and Wr+Vr/parental mean graph (b) for reducing sugar

**Figure 9:**
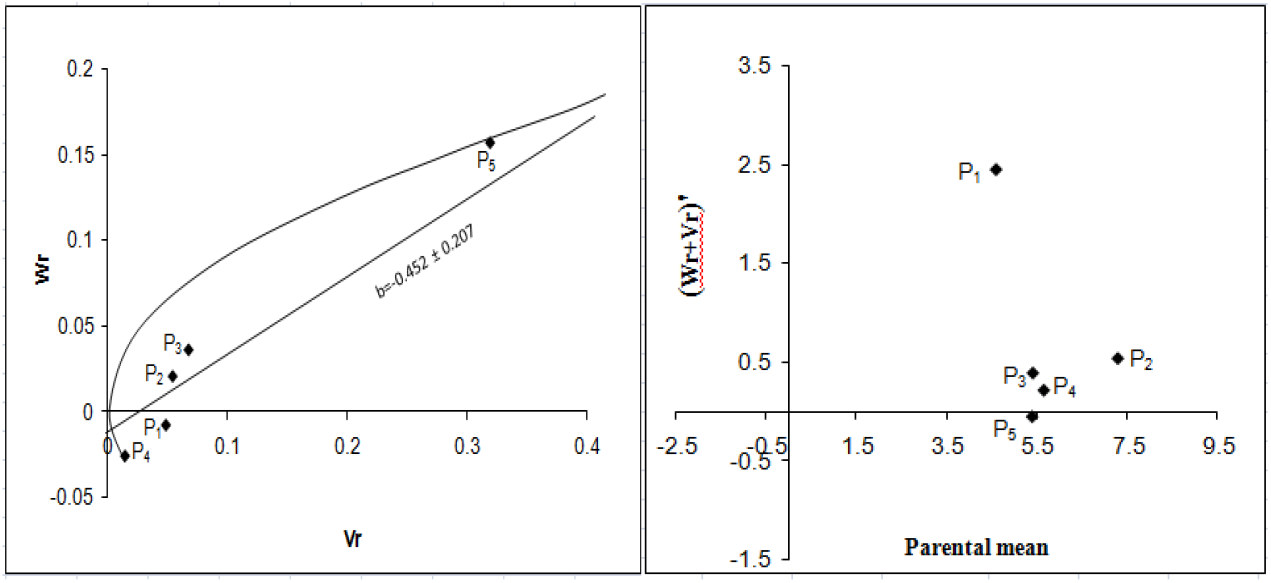
Vr - Wr graphs (a) and Wr+Vr/parental mean graph (b) for non-reducing sugar

**Figure 10:**
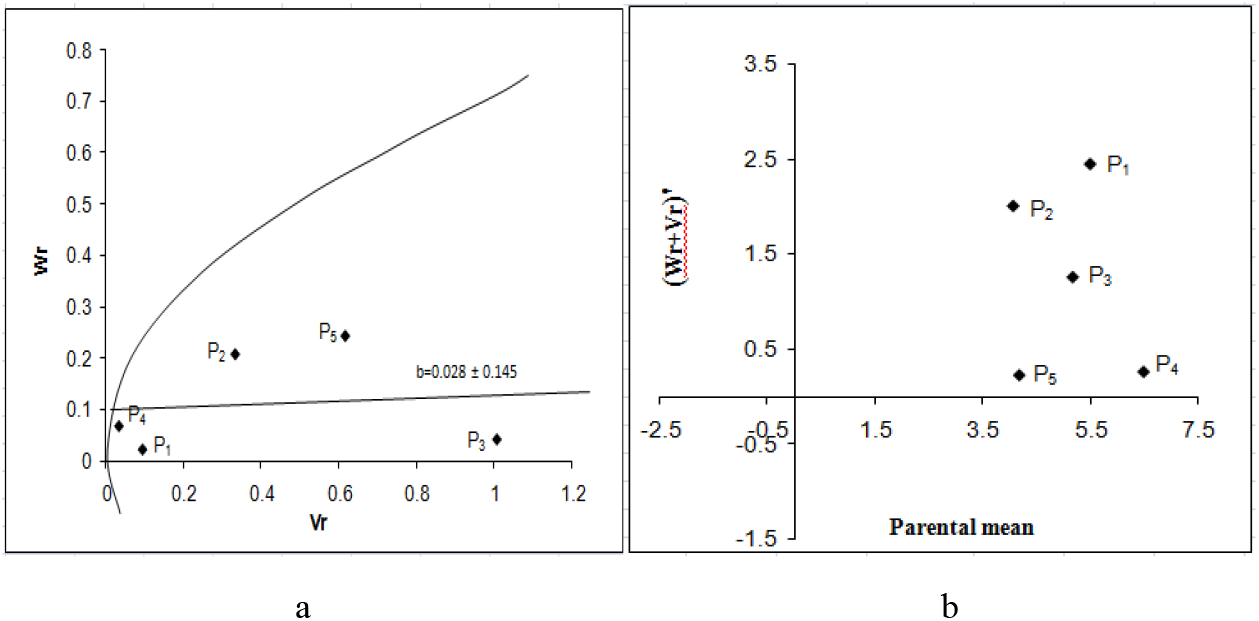
Vr - Wr graphs (a) and Wr+Vr/parental mean graph (b) for total sugar

### Reducing sugar

The Vr - Wr graph (Fig. 8a) for reducing sugar showed that the regression line passed above the point of origin which indicated the presence of partial dominance for reducing sugar. The distribution of array points in the graph suggests that P_4_ had maximum frequency of dominant alleles, P2 and P_5_ apparently contained frequency of dominant alleles while P_3_ had maximum frequency of recessive alleles as it fall far away from the origin. The regression of Wr on Vr gives the slope b = 0.075±0.104 which is less than 1, affirming presence of additive-additive nature of genetic system. Diffused array points all along the regression line in this graph indicated closer genetic diversity among parents.

The Wr+Vr/parental mean graph (Fig. 8 b) revealed that all of the parents contained the recessive alleles. The parent P_3_were high scoring P_5_ was low scoring and other parents were moderate.

### Non reducing sugar

The regression of Wr on Vr for non reducing sugar (Fig. 9a) gave a slope b = -0.452 ± 0.207 which was significantly far away from 1.0 indicating presence of additive-dominance nature of genetic system. The regression line intersected below the point of origin suggested over dominance gene action for controlling the trait. The distribution of array points indicated that among five parents, P_4_ contained the maximum frequency of dominant alleles as it held the closest position to the point of origin and P_1_ and P_2_ apparently contained frequency of dominant alleles. The parent P_5_ contained the maximum frequency of recessive alleles. Array points sprinkled all along the regression line in this graph indicated genetic diversity among parents.

The Wr+Vr/parental mean graph (Fig. 9b) tested the consistency of dominance against parental score. Parental mean suggested that all the parents contained the recessive alleles. Among them P_5_ had the highest value as it held on the top most position. So, higher non reducing sugar was associated with recessive alleles in the direction of higher values.

### Total sugar

The regression of Wr on Vr for total sugar (Fig. 10a) gave a slope b = 0.028 ± 0.145 which was less than 1.0 indicating presence of additive-additive nature of genetic system. The regression line intersected above the point of origin suggesting partial dominance gene action for controlling the trait. The distribution of array points indicated that among five parents, P_4_ and P_1_ contained the maximum frequency of dominant alleles. The maximum frequency of recessive alleles was found in P_3_ as it falls far away to the point of origin.

The Wr+Vr/parental mean graph (Fig. 10b) tested the consistency of dominance against parental score. All the five parents possessed recessive alleles associated with positive effect for total sugar content.

### Fruit yield

The regression of Wr on Vr for fruit yield (Fig. 11a) gave a slope b = 0.145 ± 0.189 which was less than 1.0 indicating presence of additive-additive nature of genetic system. The regression line intersected below the point of origin suggesting over dominance gene action for controlling the trait. The distribution of array points of five parents indicated that P_3_ and P_4_ contained the maximum frequency of dominant alleles. The maximum frequency of recessive alleles was found in P_5_ as it falls far away to the point of origin. Dispersed array points all along the regression line in this graph indicated genetic diversity among parents.

**Figure 11:**
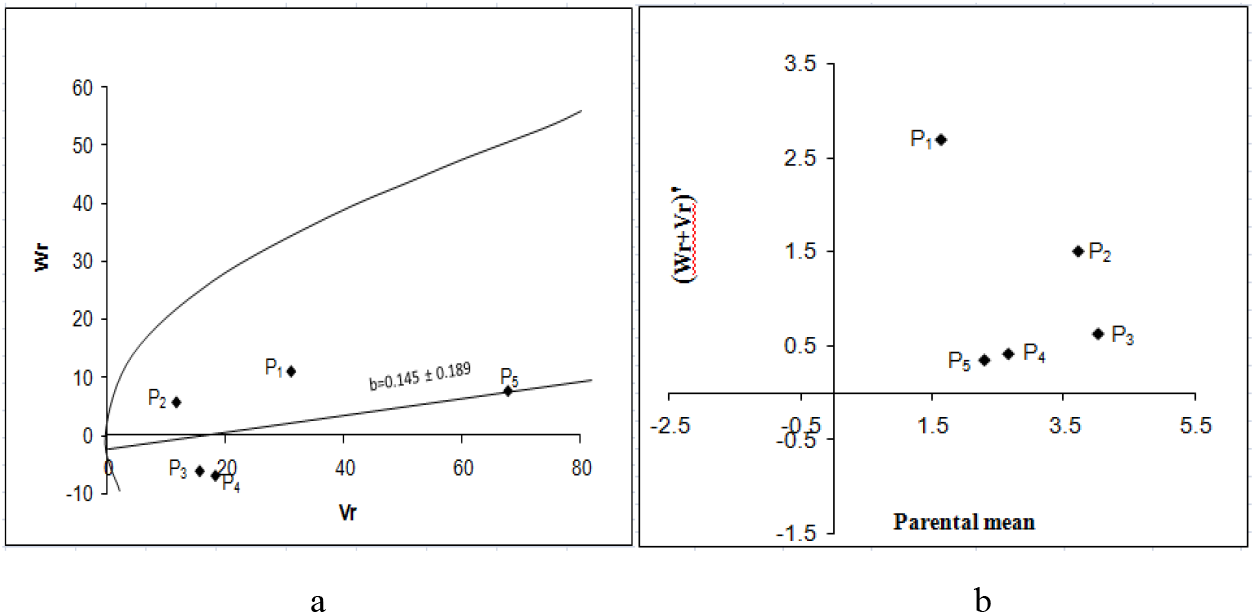
Vr - Wr graphs (a) and Wr+Vr/parental mean graph (b) for fruit yield

The Wr+Vr versus parental mean graph confirmed that all the five parents possessed recessive alleles associated with positive effect i.e. high fruit yield (Fig. 11b).

## Conclusions

The present experiment was undertaken to study the heterosis and combining ability of different quality traits of pumpkin. Eleven quality characters viz. **fruit length, fruit breadth, hollowness, flesh thickness, dry matter, brix (%), reducing sugar, non-reducing sugar, total sugar and fruit yield** were noted in a 5 × 5 full diallel population. Twenty hybrids were evaluated along with five parents to assess the heterosis and combining ability of parents for different quality characters.

Analysis of variance showed significant variation among the genotypes for fruit length, fruit breadth, hollowness, brix and fruit yield. General combining ability variances were significant for fruit length, fruit breadth, hollowness, dry matter and **brix (%)**. Specific combining ability variances were significant for fruit length, fruit breadth, hollowness, **brix (%)** and fruit yield. Reciprocal effect was found significant for fruit length, fruit breadth, hollowness and **brix (%)**. The studies on heterosis and combining ability revealed that the GCA variance estimates were found higher for five characters viz., fruit length, hollowness, dry matter, β carotene and reducing sugar indicating predominance of additive gene action. The estimates of SCA variance were high for fruit breadth, flesh thickness, **brix (%)**, non reducing sugar, total sugar and fruit yield indicating predominance of non-additive gene action in expression of these traits. The estimates of GCA effects showed no single parent contained all of the desirable characteristics. The parent P_1_ (IBD23) was good combiner for fruit length. Other good general combiners for different characters were: P_3_ (IBD47) for non reducing sugar, P_5_ (IBD57) for fruit length and dry matter content. Except these three (fruit length, dry matter content and non reducing sugar) characters, good combiner parent for other eight characters was not found.

The study of SCA effect revealed that in most of the cases combination of good x poor or even poor x poor crosses exhibited high SCA effects for many characters rather than good x good cross combinations indicating the importance of gene interactions. The best specific combiners were IBD40 X IBD47 for beta carotene, total sugar and fruit yield; IBD23 X IBD40 for brix (%), hollowness and flesh thickness; IBD40 X IBD57 for fruit breadth; IBD47 X IBD50 for non reducing sugar; and IBD47 X IBD57 for reducing sugar.

Profound reciprocal effect i.e. maternal effect was observed in most of the crosses. The cross combination IBD40 X IBD47 had positive and significant reciprocal effect for seven characters including fruit yield. Thus this combination can be a combiner for fruit yield. Reciprocal effect was also found for other traits viz. the combination IBD23 X IBD40 had positive reciprocal effect in five traits (hollowness, dry matter content, beta carotene, reducing and non reducing sugar) and two negative reciprocal effect (fruit yield and fruit length); IBD40 X IBD57 for **brix (%), reducing sugar, total sugar;** IBD47 X IBD57 for total sugar, dry matter, fruit breadth and IBD47 X IBD50 for flesh thickness dry matter, fruit breadth, beta carotene and all sugar traits.

The Vr-Wr graphs exhibited complete, partial and over dominance effect of genes for different characters. Complete dominance was observed only for beta carotene and over dominance was found for hollowness and flesh thickness. Partial dominance was observed for fruit breadth, dry matter, **brix (%)**, reducing sugar, non reducing sugar, total sugar and fruit yield.

Significant heterosis of some crosses against mid and better parent was observed for some characters. Significant highest average heterosis (mid parent) was observed for fruit yield and brix by IBD40 X IBD47, for flesh thickness, dry matter and beta carotene by IBD23 X IBD40, for reducing sugar by IBD23 X IBD50 while significant highest heterobeltiosis (better parent) was observed for fruit yield and total sugar by IBD40 X IBD47, for fruit leanth by IBD40 X IBD57.

